# Declining old pole physiology gradually enhances gene expression asymmetry in bacteria

**DOI:** 10.1101/2023.08.07.552301

**Authors:** Audrey M. Proenca, Murat Tuğrul, Arpita Nath, Ulrich K. Steiner

**Affiliations:** Freie Universität Berlin, Institute of Biology, Evolutionary Demography group, Königin-Luise-Str. 1-3, 14195 Berlin, Germany

## Abstract

Gene expression is a heterogeneous process at the single-cell level. This heterogeneity is often coupled to individual growth rates, which are also highly stochastic, leading to the emergence of multiple physiological states within bacterial populations. Although recent advances have shown that cellular aging acts as a deterministic driver of growth asymmetry, the relationship between aging and gene expression heterogeneity remains elusive. Here we show that old poles undergo a progressive decline in gene expression as mother cells age, contributing to enhance phenotypic heterogeneity in bacterial populations. We quantified the activity of promoters with distinct activity profiles: a constitutive promoter, whose expression positively correlates with growth, and the promoter of RpoS, the general stress response sigma factor, for which growth and expression are mutually inhibitory. We demonstrate that mother cells have lower gene expression for both promoters. This asymmetry could not be explained by metabolic rate differences, but rather by the increasing intracellular heterogeneity of mother cells. As a mother ages, the declining activity of its old pole produces intracellular gradients in gene expression. This intracellular asymmetry manifests in the next generation as mother-daughter asymmetry, thus representing a source of phenotypic heterogeneity for the population. Our results show that bacterial asymmetry is built into the declining physiology of mother cells across generations, illustrating the deterministic nature of aging in bacterial systems. These findings provide further evidence for cellular aging as a mechanism to enhance the variance of metabolic states found in bacterial populations, with possible consequences for stress response and survival.

## Introduction

Within isogenic populations, individual cells exhibit heterogeneity in gene expression, protein partitioning, and growth rates. This physiological variability often results from the propagation of stochastic fluctuations across regulatory networks (1, 2), leading to the coexistence of distinct metabolic states within populations (3, 4). In bacterial populations, for example, such gene expression fluctuations can lead some individuals to reproduce more quickly while others enter a slow-growing or dormant state (5). Rather than being detrimental for the population, this physiological heterogeneity is closely tied to survival in stressful conditions or changing environments (6–8). Nonetheless, recent studies have demonstrated that deterministic factors, such as cellular aging, also contribute towards the phenotypic heterogeneity previously attributed to stochasticity alone (9–11).

Aging, a physiological decline over time (12, 13), propagates along bacterial lineages across generations. According to classic theories, aging requires a clear separation between the soma and germline (14), or at least some degree of asymmetry among them (15). The fact that bacteria usually divide with morphological symmetry has, consequently, led to the assumption that these organisms should flee aging (14, 16, 17). However, the notion of symmetric fission in unicellular organisms has been overhauled in the last decades (18–21). Even in *Escherichia coli*, rod-shaped bacteria with apparent morphological symmetry, aging is determined by the asymmetric inheritance of cell poles. Bacteria must synthesize a new pole at the central plane after each fission. In the following division, one cell inherits this newly formed pole, while the other receives an old pole that can be inherited over many generations (Figure 1A). The deterministic inheritance of old poles correlates with reduced growth, lower division rates, and increased mortality, whereas the inheritance of new poles leads to higher fitness (11, 18, 19, 22). The resulting physiological heterogeneity likely drives the variation in demographic fates among cells of identical genetic background in the same environment (23). In fact, when populations experience oxidative stress, the inheritance of old poles leads to a higher probability of death, while new-pole cells continue to proliferate (11, 24). It is therefore reasonable to assume that cells receiving the old pole progressively accumulate an “aging factor”, which is asymmetrically partitioned upon division (25, 26). This process may be described as a mother cell that ages by retaining the old pole, while producing daughter cells that inherit newly synthesized poles with each division. Here we will adhere to this perspective, as it refers back to the concept of separation between soma and germline (27) and simplifies the definition of aging at the single-cell level.

**Figure 1.**
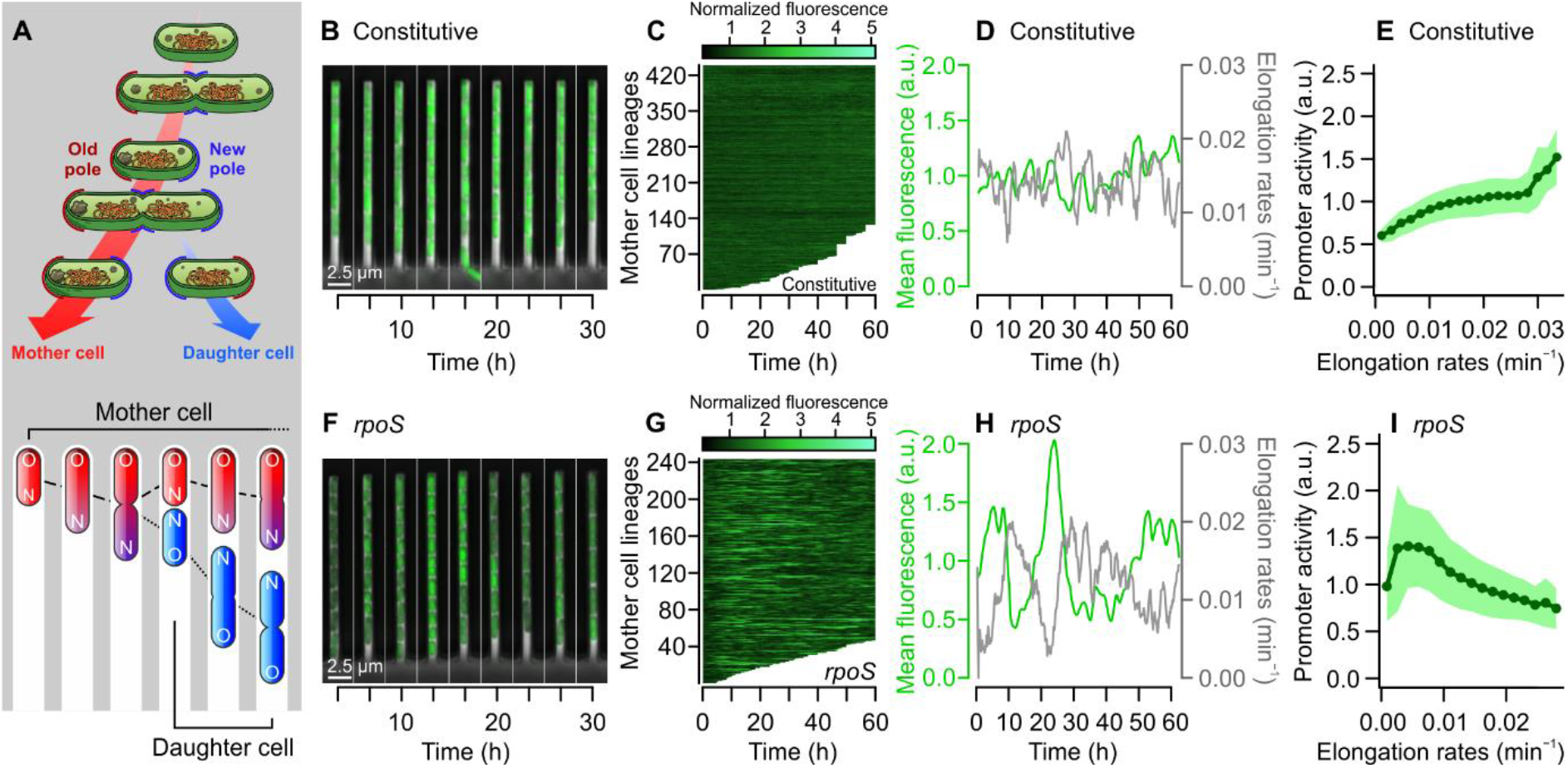
Gene expression stochasticity and correlation with growth demonstrate contrasting promoter dynamics. (A) Graphical representation of bacterial aging through asymmetric pole inheritance (top) and cells growing in the mother machine device (bottom). Old cell poles, inherited by mother cells with each division, remain trapped at the closed end of growth wells. When the mother divides, it produces a daughter cell that inherits its new pole. To investigate whether this asymmetry impacts gene expression across generations, we tracked bacteria expressing GFP from promoters with contrasting dynamics. (B to D) Mother cells constitutively expressing GFP showed little fluctuation in fluorescence levels over time. (E) Across the population (n = 62,533 cell divisions), cells exhibited a positive correlation between elongation rates and constitutive promoter activity. In contrast, (F to H) mother cells expressing GFP from the *rpoS* promoter showed highly stochastic transcriptional bursts that coincided with periods of slower growth (see also Figure S1). (I) This pattern was also present across the whole population, with slow-growing cells displaying high *rpoS* promoter activity (n = 42,350 observations). Bins = mean ± SD. Mother cells in C and G were sorted by longevity, with interrupted lines representing cells that died or were washed away.

A possible mechanism of this physiological heterogeneity is the asymmetric partitioning of intracellular damage (22, 28, 29). Misfolded proteins, for instance, tend to accumulate in the form of aggregates that eventually become too large to diffuse across the nucleoid, thus becoming trapped at the cell poles (30–32). The inheritance of protein aggregates has been shown to negatively correlate with growth rates (30, 33), but more recent studies suggest that other factors must be involved in bacterial aging. Large protein aggregates are generally absent in unstressed populations (34), and yet the physiological distinction between mothers and daughters is observed in healthy cells (10, 35). In addition, for cells under heat stress, the rejuvenation of the daughter depends on the chronological age of the mother (11), indicating that daughters can inherit intracellular damage not associated with old poles. These results suggest that, besides damage partitioning, other intracellular mechanisms also contribute to the physiological asymmetry between mother and daughter cells.

Sources of physiological heterogeneity that are considered purely stochastic can provide hints of mechanisms that influence bacterial aging (36). For example, the partitioning of freely-diffusing proteins upon division is regarded as highly stochastic process (2), but recent evidence suggests that newly synthesized gene products are partitioned with deterministic asymmetry towards daughter cells (37). It remains unclear whether this asymmetry increases as the mother cell ages, and how this pattern develops intracellularly across generations. Similarly, gene expression stochasticity propagates across regulatory networks, leading to heterogeneity at the single-cell level (2, 8, 38). It is therefore necessary to determine whether mother and daughter cells show asymmetric gene expression rates, rather than just an asymmetric inheritance of products. If divisional asymmetry develops over time and its impact can be extended to gene expression, this could reveal another mechanism of bacterial aging.

Here we address the contribution of cellular aging to phenotypic heterogeneity by investigating the asymmetry present in growth, gene expression, and protein partitioning across generations. We followed mother and daughter cells over generations, quantifying the expression and partitioning of two transcriptional reporters with opposite dynamics: one whose activity scales with growth rates, the other showing a negative correlation between growth and expression. We found that daughter cells display higher gene expression and product inheritance independently of promoter dynamics. Mother cells exhibited lower promoter activity even when elongating at a similar rate as their daughters, suggesting that gene expression asymmetry is not simply a function of asymmetric metabolic rates. We also demonstrate that the asymmetry in product partitioning originates from intracellular gradients formed within the mother cell prior to division, as an effect of old poles aging across generations. Daughter cells, whose poles are generated at a constant age, do not exhibit such decline: from birth to division, they show a homogeneous intracellular environment. Moreover, we show that mother cells compensate for a decline in old pole contribution through an increase in cell length over time. Thus, aging enhances phenotypic heterogeneity through asymmetric patterns in growth, gene expression, and product partitioning, indicating that both deterministic and stochastic factors contribute towards the variability found within bacterial populations.

## Results

### Characterization of promoter dynamics and stochasticity

To investigate the effects of aging on growth, partitioning of gene products, and gene expression heterogeneity, we imaged *E. coli* cells growing in constant exponential phase within mother machine devices (39). These microfluidic devices contain thousands of narrow growth wells connected in one end by flow channels, which supply fresh nutrients to trapped cells (Figure 1A). Throughout the experiments, the mother cell remains lodged at the closed end of the growth well, while continuously producing daughter cells. We tracked mother cells for up to 100 generations and each daughter from birth to its first division, quantifying aspects of their growth physiology (length, division intervals, elongation rates) and gene expression (mean fluorescence intensity, promoter activity).

We measured gene expression in strains containing a promoter of interest fused to fluorescent transcriptional reporters. For this, we used two promoters with contrasting dynamics: *rpoS* (40) and a constitutive promoter (41). The former expresses the RpoS sigma factor, which regulates the general stress response and the transition to stationary phase (42, 43), showing highly stochastic expression in transcriptional bursts that negatively correlate with growth rates (4, 8). The constitutive promoter, on the other hand, is expected to show a positive correlation between expression and growth (44) and lower noise levels than *rpoS* (1). By comparing the long-term activity of both reporters, we can identify effects of aging that are independent of specific promoter dynamics.

To confirm that both reporters exhibited the predicted promoter dynamics, we quantified mean fluorescence levels per mother cell lineage over time. As predicted, *rpoS* exhibited highly stochastic fluctuations and larger variance than the constitutive promoter (F test: F = 4.533, p < 0.001). This difference in noise levels can be expressed by coefficients of variation, with the constitutive reporter showing a 2-fold lower CV = 20.43% (Figure 1B to D) than *rpoS* (CV = 43.49%; Figure 1F to H). When we considered the rate of expression of each reporter (see Methods for details), the constitutive promoter showed an increase in activity with faster elongation rates (β = 13.94, t = 78.1, p < 0.001; Figure 1E). In contrast, random bursts of *rpoS* expression corresponded to periods of slower growth (Figure 1H and S1), with the relationship between elongation rates (r) and promoter activity being expressed by an exponential decay function (1.498*e^-28.898*r^, t = -80.33, p < 0.001; Figure 1I). Since few cells undergo large transcriptional bursts, *rpoS* measurements yielded long-tailed distributions of mean fluorescence and promoter activity, in contrast with the Gaussian distribution of constitutively expressed GFP (Figure S1).

As a functional control to determine that the *rpoS* reporter reflected its biological activity of this sigma factor, we induced a transition from exponential to stationary phase within the mother machine device (Figure S2). Since RpoS inhibits growth as cells enter stationary phase, we expected to find a gradual increase in mean fluorescence over time. Our results confirmed this prediction, with mother cells showing an increase in RpoS expression as their elongation rates declined (Figure S2B). This indicates that our fluorescence measurements reflect RpoS activity, with a negative correlation between growth and expression observed both during exponential phase transcriptional bursts and in the transition to stationary phase.

These results confirm the expectations for both promoters, summarizing the contrasting dynamics of *rpoS* and constitutive reporters in terms of noise and growth-dependence. The next section will explore how divisional asymmetry contributes to the observed single-cell variance in product inheritance and gene expression.

### Asymmetric product partitioning is independent of expression dynamics

The asymmetric metabolic rates of *E. coli* can be represented through a quantification of elongation rates over each generation (Figure 2A). Daughter cells displayed faster growth than mother cells (*rpoS*: t = 132.95, p < 0.001; constitutive: t = 174.42, p < 0.001), with their generation times being, on average, 1.81 min shorter (t = 32.47, p < 0.001). We can expect this asymmetry to be reflected also in gene expression. For example, previous results have shown that gene products are inherited with deterministic asymmetry (37), where daughter cells receive a higher load of newly synthesized components. It is unclear, however, whether this pattern remains true when expression is negatively correlated with metabolic rates, as it the case of the *rpoS* reporter.

**Figure 2.**
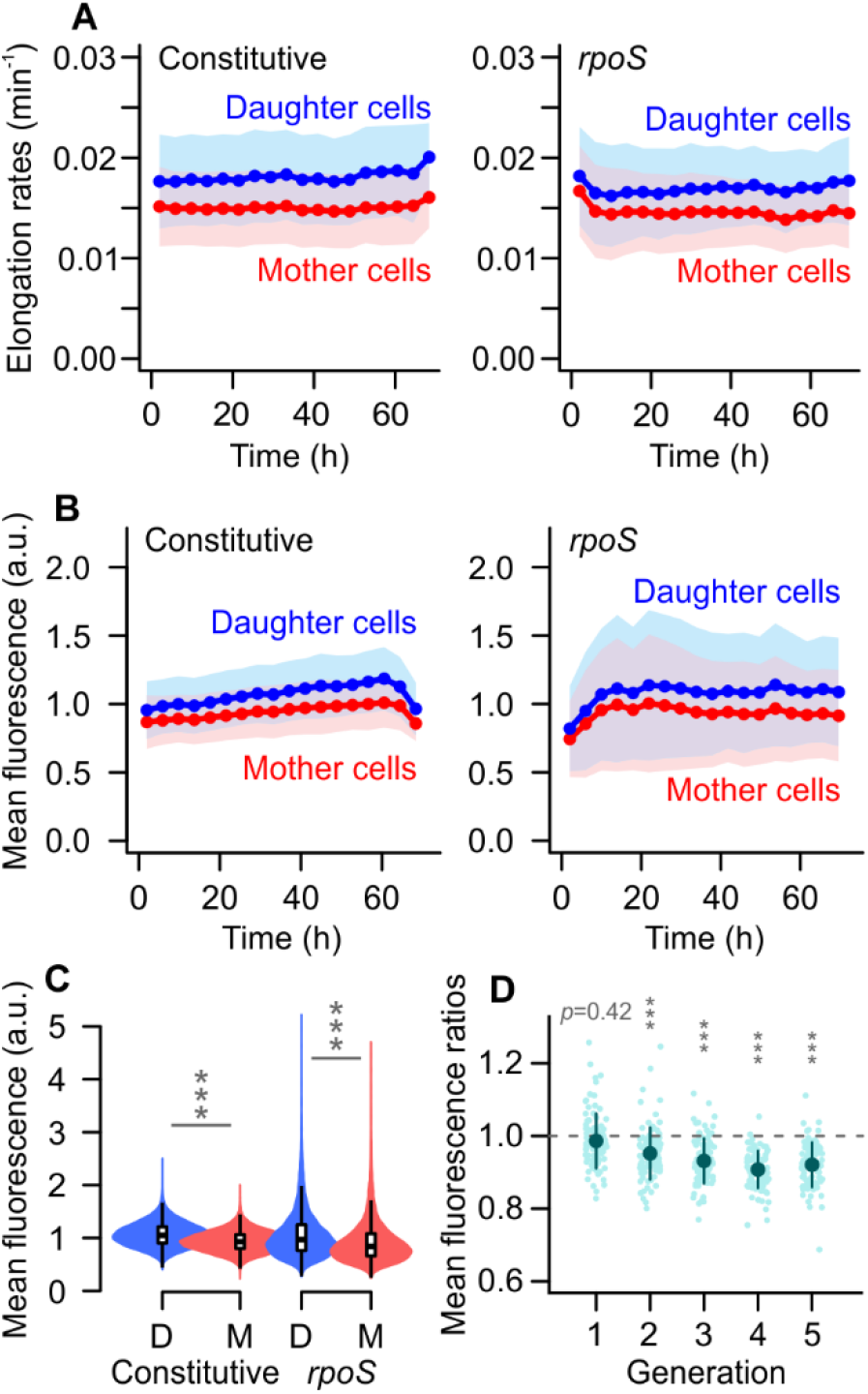
Mother and daughter cells show asymmetry in growth and gene product inheritance. (A) Daughter cells displayed faster growth, with a stable asymmetry in elongation rates over time. (B and C) Independently of the underlying promoter dynamics shown in Figure 1, mother cells exhibited lower mean fluorescence levels after division. This supports prior reports that *E. coli* partition “good” components asymmetrically upon division (37, 45). (D) To verify that this observation was not an artifact of cell positioning within the mother machine device, we measured *rpoS* fluorescence ratios between the 1st and 2nd cell in the growth well. Since the initial cell to arrive in the well can have any polarity (see diagram on Figure S3), the 1st cell has equal probability of being a mother or a daughter upon division. As such, no asymmetry is present on generation 1. From generation 2 onwards, the 1st cell inherits the old pole after each division, thus being called a mother cell. Asymmetry is then observed for every generation, showing that this pattern is age-dependent rather than a positioning artifact.

To verify this, we measured fluorescence asymmetry shortly after each division. Daughter cells showed higher mean fluorescence levels for both the constitutive (t = 189.21, p < 0.001) and the *rpoS* reporter (p = 272.76, p < 0.001; Figure 2B and C), with a slight increase in asymmetric inheritance of gene products across time (Figure S3). While the variance in elongation rates accounted for most of the deterministic heterogeneity of *rpoS* fluorescence (V_r_ = 23.5%; Table S1), a pattern that was not verified for constitutive GFP expression (V_r_ = 1.6%), divisional asymmetry had a similar contribution to the total variance in fluorescence levels for both strains (V_asym_ = 9.9% and 11.8%, respectively). Although the partitioning of gene products is a highly stochastic process, with unexplained variance accounting for 54.1% of the total heterogeneity for both reporters, these results suggest that deterministic asymmetry contributes to increase the total heterogeneity of a population.

We performed additional controls to ensure that this asymmetry was not an artifact of the mother machine architecture (SI Text). Within the device, mother cells are always lodged at the end of a growth well, whereas their daughters are in the middle (Figure S3). This means that the mother has only one neighbor, while the daughter has two. It is possible, therefore, that the light scattered from neighboring cells would make the daughter appear brighter. To investigate this, we quantified *rpoS* asymmetry after the first division recorded within the device. Because the polarity of the “founder” cell is unknown, the cell at the closed end has a 50% probability to inherit either a new or old pole. If the asymmetry observed in Figure 2B has a physiological origin, it should be absent at this first division. As shown in Figure 2D (see Figure S3 for diagrams and distributions), asymmetric product inheritance was not verified in the first division, and gradually increased in subsequent generations, indicating that asymmetric product inheritance is a consequence of bacterial aging.

### Aging contributes to gene expression heterogeneity

We next investigated whether aging is a source of gene expression heterogeneity in bacterial populations. To verify this, we calculated the rate of GFP production from each promoter, since it takes into consideration differences in individual growth rates (4, 8). This correction is essential because daughter cells elongate faster and have a larger ribosomal content than mother cells (45). Nonetheless, even after accounting for the underlying asymmetry in metabolic rates, we found that daughter cells had larger rates of constitutive fluorescence expression (Figure 3A; t = 185.14, p < 0.001). As for *rpoS* expression, we expected to find higher promoter activity among mother cells, since RpoS has growth-inhibitory activity. Instead, we verified that daughter cells also had greater *rpoS* expression (Figure 3A; t = 209.72, p < 0.001). Moreover, mothers and daughters had asymmetric gene expression even when growing at similar rates (Figure 3B). Constitutive promoter activity increased with growth (t = 30.25, p < 0.001) with a similar slope for both mother and daughter cells (β = 1.94, p = 0.084), but daughters showed higher expression rates (t = 17.47, p < 0.001). The *rpoS* promoter exhibited similar asymmetric activity (t = 23.21, p < 0.001), with a steeper negative correlation between growth and expression observed among mother cells (β = -49.56, p < 0.001) than daughter cells (β = -46.53, p < 0.001). Thus, gene expression asymmetry cannot be explained solely by a difference in metabolic rates, suggesting that other intracellular processes associated with aging might contribute to this deterministic heterogeneity.

**Figure 3.**
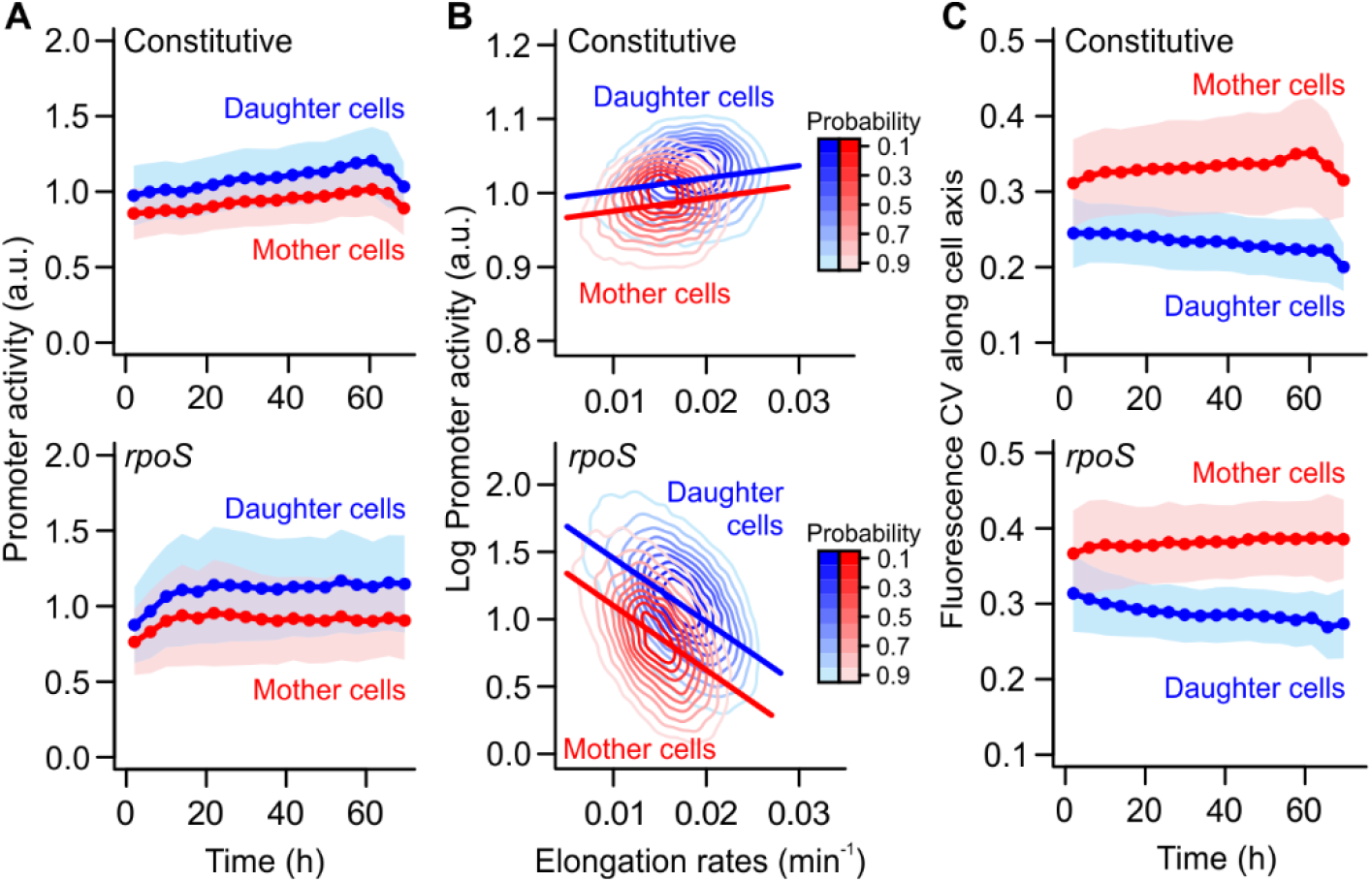
Aging leads to gene expression asymmetry independently of promoter dynamics. (A) Daughter cells exhibited higher promoter activity, whether the overall activity level remained constant or increased over time. This indicates that, besides simply inheriting more newly synthesized products, daughter cells have higher gene expression rates. (B) Gene expression asymmetry was present independently of its correlation with elongation rates. This correlation was better represented by separate regression lines for mothers and daughters, suggesting that even cells growing at a similar rate show asymmetric gene expression. Thus, the difference in promoter activity cannot be explained solely by the difference in metabolic rates shown in Figure 2A. (C) Examining the intracellular distribution of fluorescence levels, mothers and daughters also differed in the variance of fluorescence signals. (A and C) Bins = mean ± SD.

Investigating potential sources of asymmetry at the intracellular level, we asked whether mother and daughter cells differed in the intracellular distribution of synthesized products. For this, we quantified the variance in synthesized products across the length of each cell (*i*.*e*., from old to new pole). Independently of the reporter, we observed mothers harbored much larger intracellular variation than their daughters (Figure 3C; p < 0.001). This distinction increased with maternal age, with the CV of mother cells increasing over time (t_Const_ = 23.92, t_*rpoS*_ = 13.21; p < 0.001), while the fluorescence variance within daughter cells decreased (t_Const_ = -33.99, t_*rpoS*_ = -34.55; p < 0.001). This difference in intracellular variance might help elucidate the sources of asymmetry in gene expression and will be further explored in the following section.

### Mother-daughter asymmetry is built within the mother cell

To investigate the intracellular variance in gene expression, we quantified the mean fluorescence signal along the cell axis of each individual. Cells were normalized by length to allow for a comparison of these distributions, and average fluorescence transects were calculated for each generation (Figure 4). To avoid biases introduced by positioning artifacts, we performed control analyses and corrections described in SI Text and Figure S5. The resulting transects indicated that the intracellular variance found in mother cells derives from a gradient in gene expression between new and old poles. We evaluated these gradients through generalized additive models, considering the distance from each pole towards the center and the time of each measurement (in generations) as smooth terms (see Figure S6, Table S2 and S3). For both promoters, our analysis indicated that the effect of distance from the pole varied smoothly with time (p < 0.001). Additionally, expression levels in each half of the mother cell varied differently over time (p < 0.001) and across the cell length (p < 0.001). Together, these factors explained 51.3% of the maternal deviance in *rpoS* expression, and 67.1% for the constitutive reporter. Daughter cells exhibited a homogeneous distribution of intracellular fluorescence, with slightly lower fluorescence levels around the old pole area of raw transects (Figure S6 and Table S2). Although our statistical models indicated a significant difference between new and old areas of the cell (p < 0.001), each varying smoothly across length (p < 0.001) and over time (p < 0.001), these factors explained a lower fraction of gene expression deviance in daughter cells (*rpoS*: 11.7%, Constitutive: 33.3%).

**Figure 4.**
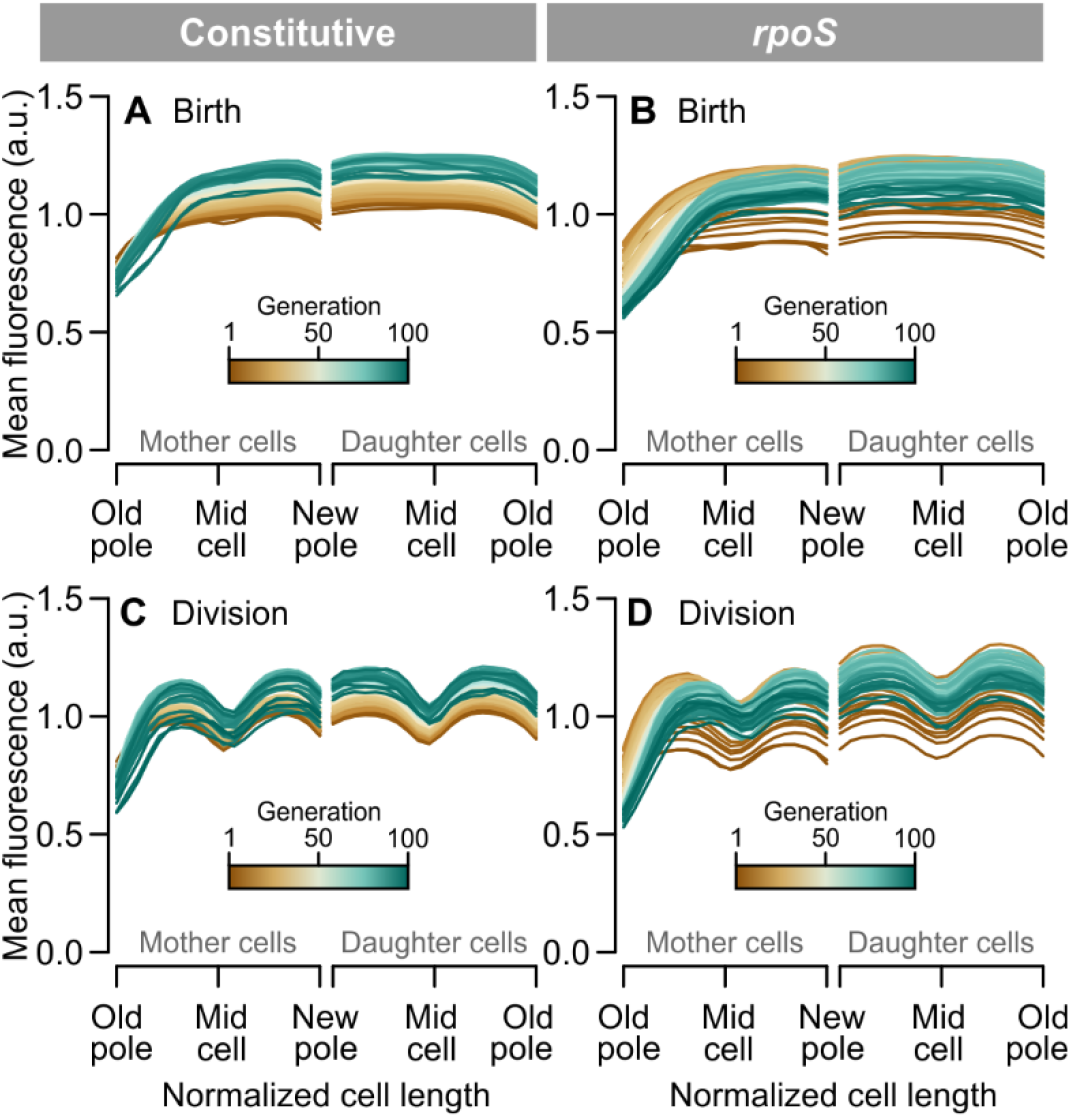
Gene expression asymmetry across the cell length. Each line represents a series of fluorescence measurements from pole to pole, averaged for all cells of a given generation. For both promoters, mother cells had darker areas around the old poles from birth (A and B) to division (C and D), indicating a gradient in gene expression along the cell length. Daughter cells, on the other hand, showed higher intracellular homogeneity in gene expression. The fission site was visible in the transects measured before division. Control analyses and transformations (length cropping) to avoid positioning artifacts are detailed in SI Text and Figure S6. These results suggest that the asymmetry between mother and daughter after division (Figure 2B) derives from the intracellular gradient built within the mother cell over time (see also Figure S7).

This intracellular asymmetry persisted throughout each cell cycle (Figure 3C and D), with the fission plane becoming visible halfway through the cell cycle. The intracellular asymmetry that an individual exhibited prior to division was correlated with the asymmetry among its offspring (Figure S6), indicating that this pattern propagates across generations. This suggests that the asymmetry in product inheritance, shown in Figure 2, derives from the intracellular asymmetry that progressively develops within mother cells across generations, with aging impacting the contribution of old poles to total gene expression.

### Pole age determines intracellular gene expression asymmetry

Since mother cells exhibit an intracellular gradient and product inheritance asymmetry increases over time (Figure S3), we hypothesize that maternal intracellular heterogeneity is age dependent. In rod-shaped bacteria, age is a function of how many times a cell pole was consecutively inherited (Figure 5A). While maternal old poles exist for up to 100 generations in our experiments (0 < age ≤ 100), their new poles are always synthesized at the last fission event (age = 0). The new pole of the daughter cell, consequently, also has age = 0, while its old pole has age = 1. As such, we expect old poles of mother cells to be physiologically older, while new poles of mothers and daughters should be equivalent. This is demonstrated through fluorescence measurements in Figure 5. For both reporters, maternal old poles exhibited lower accumulation of gene products than old poles of daughter cells (Figure 5B; t_Const_ = 77.50, t_*rpoS*_ = 50.79, p < 0.001). Fluorescence levels in new poles, on the other hand, showed little difference between mothers and daughters (Figure 5C; t_Const_ = 22.57, t_*rpoS*_ = 10.60, p < 0.001). The greater homogeneity of daughter cells can also be visualized through the ratios between new and old poles. Daughter intracellular ratios approached 1 (Figure 5D), showing little change over time (*r*_*rpoS*_ *=* 0.09, *r*_Const_ = 0.02), since the age of both poles remains constant for daughter cells. Mother cells, in contrast, exhibited increasing intracellular asymmetry with each division (*r*_*rpoS*_ *= -*0.57, *r*_Const_ = - 0.26), which coincides with the increasing age difference between maternal old and new poles.

**Figure 5.**
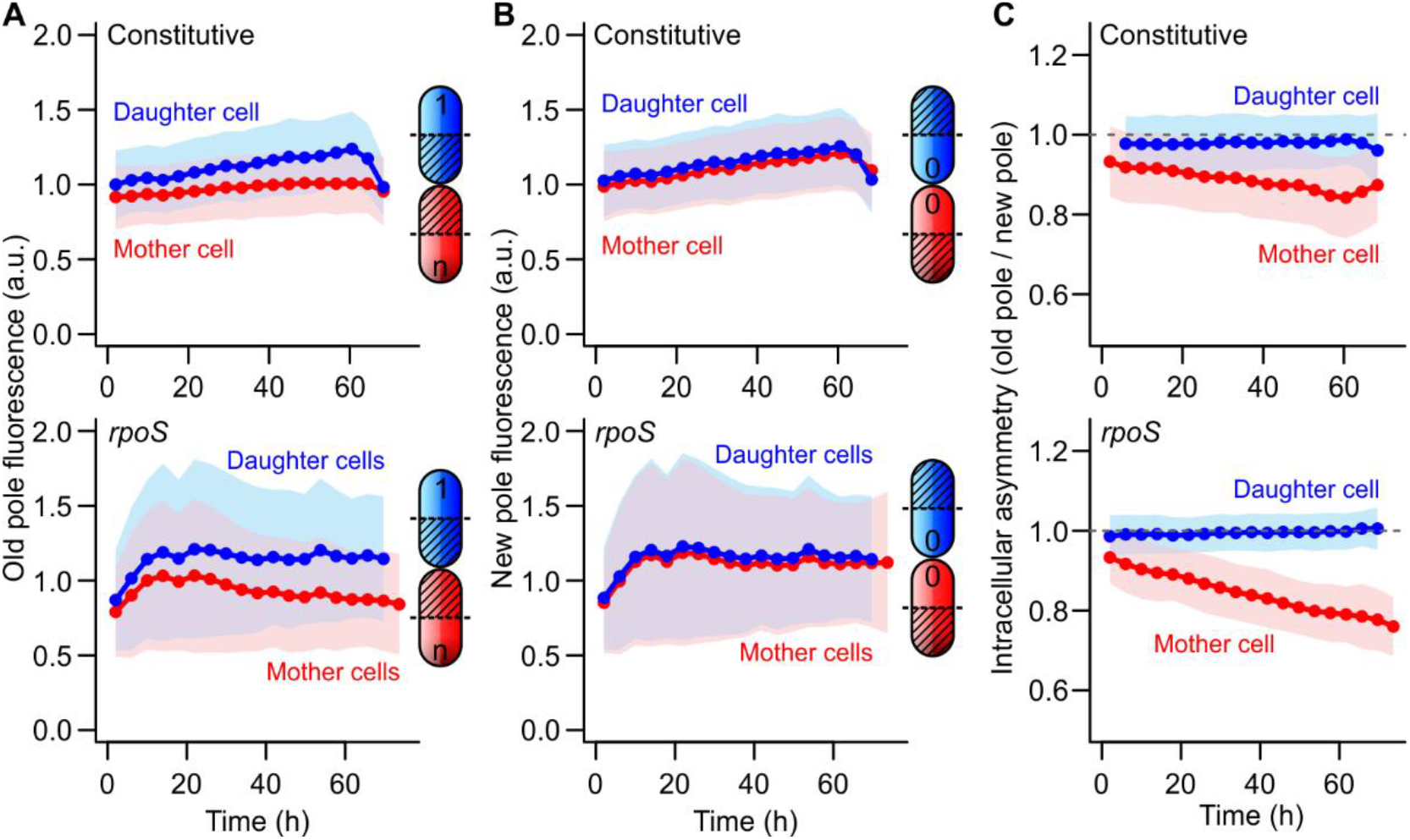
Intracellular asymmetry depends on the age of cell poles. For lineages in a mother machine device, maternal old poles become older with each generation (1 < n < 100), while old poles of daughter cells always have age = 1. This difference in pole ages predicts that most of the difference between mother and daughter cells should come from their old poles. (A) Fulfilling this prediction, we verified that old poles of daughter cells exhibited higher fluorescence expression than maternal old poles. (B) In contrast, new poles of mother and daughter cells showed little difference in gene expression, since both new poles are generated at the fission site. (C) The age difference between new and old poles increases over time for mother cells, leading to an increase in intracellular asymmetry as the mother ages. Daughter cells remain nearly symmetric over time, coinciding with a constant age difference between poles. Bins = mean ± SD.

To determine the effects of bearing old poles on maternal gene expression, we calculated the length of these poles based on *rpoS* expression gradients. For each cell, we performed a segmented regression analysis to find the breakpoint that determined the transition from the polar gradient to the mid-cell fluorescence plateau. The distance from the cell pole to this breakpoint represents the old pole length, shown in Figure 6A. Both the old pole length (t = 84.44, p < 0.001) and the overall cell length (t = 47.37, p < 0.001) increased over time, though at distinct rates (ANCOVA interaction: F = 87.08, p < 0.001). Combining these measurements, we observed an increase in the fraction of the mother cell represented by the old pole (t = 50.82, p < 0.001), from 25.3% to 36.2% after 72 h (Figure 6B). Because of the increasing maternal length, a morphological asymmetry between mothers and daughters developed over time (Figure 6C, β = 0.002, t = 51.32, p < 0.001). This indicates that the progressive change in maternal length derives from off-center divisions, which are also visible in Figure 4CD. To investigate whether these changes could explain gene expression asymmetry at the individual level, we recalculated *rpoS* activity estimates while excluding the old pole area of mother cells (Figure 6D and E). Both mean fluorescence (t = 28.31, p < 0.001) and promoter activity (t = 164.58, p < 0.001) increased, with the former approaching daughter cell levels. The fact that promoter activity remained asymmetric despite the removal of old poles indicates that other factors, such as metabolic rates, remain an essential component of bacterial asymmetry.

**Figure 6.**
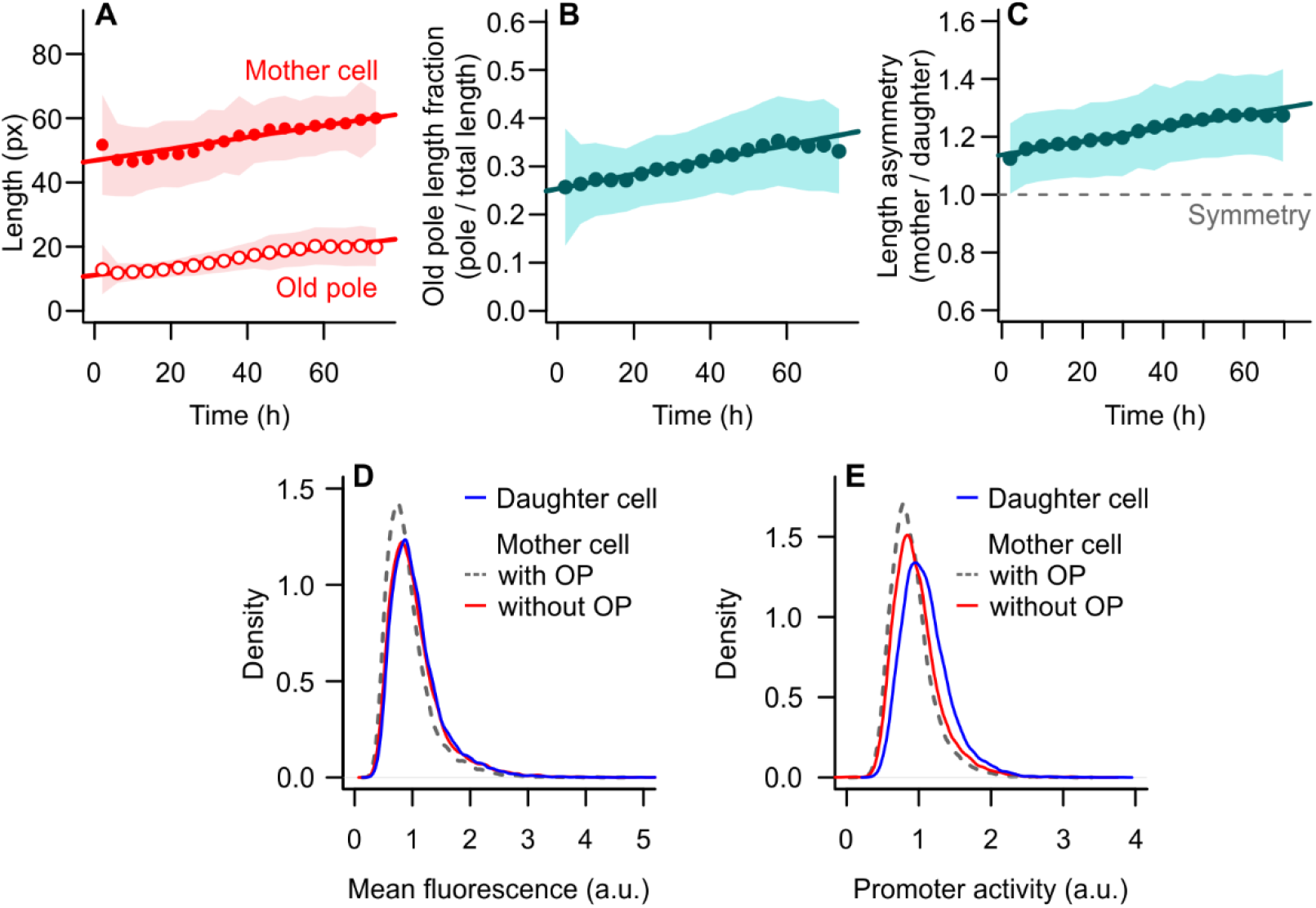
Lengthening of the old pole contributes to decline in gene expression. (A) Mother cells increase in birth length over time, which is accompanied by an elongation of the old pole area of the cell. (B) With age, old poles represent an increasingly larger proportion of the mother cell. (C) Since daughter cells carry old poles of a constant age, the morphological asymmetry between mothers and daughters increases over time. This suggests that aging in *E. coli* is not just a matter of physiological asymmetry, but also comprises a morphological asymmetry that develops as the mother ages. (A to C) Bins = mean ± SD. (D and E) If we remove the old pole are of the mother cells and re-estimate mean fluorescence and *rpoS* promoter activity, these measurements approach the levels of daughter cells. This suggests that bearing old poles contributes to the lower gene expression observed in mother cells.

Taken together, these results indicate that increasingly aged old poles promote a gradual decline in maternal gene expression, producing intracellular asymmetry gradients that are propagated across generations. Through divisional asymmetry, bacterial aging contributes to enhance not only the heterogeneity in *E. coli* growth rates, but also in gene product inheritance, gene expression, and cell length.

## Discussion

Gene expression, just like growth, is highly stochastic in bacteria. Nonetheless, we demonstrated that part of the variation commonly attributed to stochastic factors is due to cellular aging. Mother cells exhibited lower gene expression rates than their daughters (Figures 2 and 3), independently of whether the activity of a given promoter was positively or negatively correlated with growth. Since this asymmetry was present even when mothers and daughters elongated at similar rates (Figure 3B), this phenomenon cannot be explained by the higher metabolic rates of daughter cells, but rather by cellular aging. Across generations, an asymmetry gradient developed within the mother cell due to the age difference between its old and new poles (Figures 4 and 5), which manifested in the next generation as the asymmetry between mother and daughter. The daughter cell, which has poles of similar age, exhibited homogeneous gene expression. Finally, we showed that mother cells bear an increasingly large old pole area as they age, suggesting that the progression of cellular aging leads to morphologically asymmetric divisions in *E. coli*. If this old pole area were to be removed, mothers and daughters would become more symmetric (Figure 6). Taken together, these results indicate that aging processes contribute to increasing the variance in gene expression within bacterial populations.

A comparable paradigm shift occurred when previous studies demonstrated the influence of cellular aging over the distribution of bacterial growth states. Although bacteria grow at a constant rate when maintained in stable environmental conditions (39, 46), mother and daughter cells reach distinct states of growth stability (10), building on the notion that the inheritance of old cell poles leads to lower fitness (18, 19, 30). This deterministic asymmetry can be advantageous over the purely stochastic partitioning of damaged components (9, 26). Nonetheless, stochastic components have a core relevance for bacterial physiology, dictating mortality patterns of mother and daughter cells (11), and propagating through genetic regulatory networks (1, 2). One example of a highly stochastic component is the sigma factor RpoS, whose pulsating dynamics of growth inhibition can lead to higher stress survival (4, 8, 47). By showing that mother and daughter cells have asymmetric *rpoS* expression, our results suggest yet another layer to the phenotypic heterogeneity modulated by aging. It remains an open question whether the higher levels of RpoS observed in daughter cells could also be involved in their higher probability of survival to oxidative stresses (24).

The evidence that asymmetry is also present at the gene expression level also invites a new mechanistic perspective on bacterial aging that links molecular to organismal processes. For instance, Shi *et al*. demonstrated that daughter cells receive more newly synthesized proteins upon division (37), although this study was limited to a few generations. Our results suggest that this partitioning asymmetry does not rely on the underlying dynamics of promoter activity, and that it develops over time as a consequence of the physiological decline observed in maternal old poles. Moreover, we show that bacterial asymmetry extends beyond the simple inheritance of genetic products, with daughter cells exhibiting higher rates of gene expression. This can be partly explained by the fact that ribosomes are also partitioned asymmetrically, with daughter cells inheriting a larger fraction (45). However, while ribosome contents are highly correlated with elongation rates (48), gene expression asymmetry was present even after correcting for differences in growth. Similarly, we demonstrated that maternal old poles exhibited a continuous physiological decline despite elongation rates remaining constant. It is likely that a combination of factors, from the inheritance of newly synthesized components to the accumulation of damaged proteins, interacts to produce the progressive decline we report.

Although traditionally the maternal aging phenotype has been mostly attributed to the accumulation of misfolded proteins (10, 30, 32, 33), recent studies have suggested that the inheritance of protein aggregates does not correlate with slower elongation rates (34, 35). It is important noting, nonetheless, that whether or not the accumulation of protein aggregates in the old poles is correlated with a decline in growth, it could impact other physiological processes. For instance, the presence of aggregates can displace the FtsZ ring and generate morphological asymmetry upon division (49), which would explain the increase in maternal length with age described in Figure 6. We do not discard the possibility that such aggregates might displace newly synthesized proteins and ribosomes, contributing to produce intracellular gradients in mother cells. Further studies are necessary to investigate the correlation between protein aggregation, growth, and gene expression across generations.

We have shown that cellular aging contributes to the phenotypic heterogeneity found within bacterial populations, influencing growth and gene expression. More than producing a fixed asymmetry between mothers and daughters, the progressive aging of old poles widens the physiological heterogeneity with each division. By linking gene expression and aging dynamics, our findings provide a framework for future studies on the causal understanding of bacterial aging. Moreover, our results invite the question of which other mechanisms of metabolic regulation might be involved in the aging process. It is possible that the maturation and physiological decline of the old poles have direct consequences for bacterial growth — *e*.*g*., the old area, representing a considerable fraction of the mother cell, might have a lower contribution to cell wall elongation, producing differences in elongation rates among mother and daughter cells. Taken together, our findings demonstrate that cellular aging is an essential part of non-genetic variance in bacterial populations, and future studies might elucidate its role as a driver of phenotypic heterogeneity involved with survival to oxidative stress and antibiotic treatments.

## Methods

### Strains and growth conditions

*E. coli* K-12 strain MG1655 containing a plasmid-based GFP reporter of RpoS expression was obtained from the U. Alon library (40). Extensive characterizations of reporters in this library, including of RpoS and downstream promoters, demonstrated that activity patterns observed on a plasmidial reporter do not differ from a chromosomally integrated one, except for a lower fluorescence intensity of the latter (4, 8, 38). Reporters from this library have also been shown to match biological expression levels (50). Since imaging strains with a chromosomal integration would require higher exposure to phototoxic stress, we used the original strain to minimize damage exposure. RpoS expression patterns were compared with constitutive GFP expressed under the PA1 promoter (41). Liquid cultures were performed on M9 media (1xM9 salts, 2 mM MgSO_4_, 0.1 mM CaCl_2_) supplemented with 0.2% Casamino Acids and 0.4% glucose. For microfluidic experiments, 0.075% Tween20 (Sigma-Aldrich) and 1 μg/ml Propidium Iodide (PI) were added to the media. Tween20 prevents cell adhesion to the walls of a microfluidic device, and PI is a marker of cell lysis. All experiments were conducted at 37°C.

### Microfluidic design and fabrication

The mother machine microfluidic design was used for cell capture and single-cell imaging (39). The design consisted of four parallel flow channels (1 cm x 80 μm x 10 μm), each containing 1,000 growth wells (25 x 0.8 x 1.2 μm). Each growth well contains a mother cell lineage, which receives fresh culture medium diffused from the flow channels throughout the experiment. The original mold was fabricated through laser etching, and epoxy copies were produced for constant handling. Individual devices were fabricated with Sylgard 184 polydimethylsiloxane (PDMS) (Dow Corning, USA) using epoxy copies as negative molds for PDMS casting. The cast was degassed in a vacuum chamber and hardened at 80°C for 1h, then removed from the mold and punctured with a 0.5 mm biopsy punch to create inlet and outlet ports. Each device was thoroughly washed with ethanol and water, dried, and attached to 24 x 40 mm coverslips through plasma bonding. A final curing step was performed by incubating bonded devices at 60°C overnight. Prior to use, devices were plasma-activated and treated with 20% polyethylene glycol (PEG) for 1 h.

### Cell loading and device setup

Overnight bacterial cultures were diluted into 15 ml M9 media and grown to exponential phase for 2 h prior to loading (until OD_600_ = 0.4-0.6 was reached). Cultures were centrifuged at 4000 rpm for 10 min, then resuspended into 200 μl of M9+Tween20+PI. These concentrated cultures were injected into the devices through the outlet ports, and devices were centrifuged at 1500 rpm for 8 min to push bacteria into the growth wells. Once the proper loading was verified, the flow media input was connected to each device through Tygon tubing (ND 100-80, Saint-Gobain, Darwin Microfluidics). A constant flow of 200 μl/h was ensured by a peristaltic pump (Ole Dich, Denmark). In the segment of the inlet tubing that ran through the pump, Tygon tubing was replaced by flexible pumping tubing (Masterflex Tygon E-3603, VWR). Outflow tubing were connected to outlet ports, running to a waste container. Loaded devices were placed in the microscopy incubation chamber at 37°C (Pecon TempController 2000-1).

For stationary phase experiments (Figure S2), this setup was combined with an inlet of spent M9 medium. An overnight culture was grown in M9, which was then centrifuged and filter-sterilized. This spent medium was supplemented with 0.075% Tween20 and 1 μg/ml PI, and connected to the regular media inlet with a replacement rate of 600 μl/h.

### Microscopy and image acquisition

Time-lapse imaging was performed on a Nikon Eclipse Ti2 inverted microscope equipped with a motorized stage and Perfect Focus System (PFS), using a 100x oil objective and DS-Qi2 camera. Automated image acquisition was controlled through the NIS Elements software, configured for phase contrast acquisition in 2 min intervals for 72 h. In order to decrease photooxidation effects (24), fluorescence images for GFP and RFP quantification were obtained in 10 min intervals.

### Image processing

Images were pre-processed on ImageJ for chromatic shift correction and background subtraction (rolling ball radius = 20 px, sliding paraboloid). Automated image processing was performed using the DeLTA deep learning-based software (51), for which the deep learning algorithm was further trained on diverse single-cell data from our experimental system. Segmentation was performed on phase contrast images, and measurements were obtained for cell length, area, division intervals, and mean fluorescence intensity. Fluorescence transects were obtained by extracting the mean fluorescence intensity of each pixel along the mid-cell line. Segmentation and tracking errors, though rare, were corrected with a custom program implemented in R v.4.1.2 (52). Manual corrections were performed when necessary. Each lineage was followed until the death of the mother cell, and only complete generations were considered for statistical analyses (*i*.*e*., daughter cells that left the growth well before a division was observed were excluded from measurements).

### Statistical analysis

Statistical analyses on the output growth and fluorescence data were performed in R v.4.1.2, considering mother cells for the entirety of the experiments and the first generation of each daughter cell. For the comparison of individuals at the same stage of the cell cycle, fluorescence measurements were taken from the first frame after division (indicated as “birth” frame on Figure 4) unless otherwise indicated. Elongation rates were calculated for each cell from an exponential fit to length measurements of each generation. As such, mean fluorescence and elongation rate estimates for mothers and daughters are paired for every division. When parametric tests on fluorescence data were necessary, the data was log-transformed. On all representations of binned data, bins represent mean ± standard deviation (SD), and bins comprising fewer than 10 measurements were excluded.

Fluorescence transects shown in Figures 4 were normalized by length for graphical representation and for analysis through generalized additive models, which were performed using R package mgcv v.1.3-38 (53). While the transects in the main figure were averaged for each generation, individual measurements were used in the analysis. We focused on the differences between new and old cell areas for the construction of the models, as shown in Table S2. Each fluorescence transect was cut in half, thus splitting the new and old areas of each cell, and the distance from the pole towards the center was used as a smoothing term in the model (see also representation in Figure S7A and B). The model structure that provided the lowest AIC was considered optimal (Table S2) (54). Transects shown in Figure 4 were cropped on both ends before length normalization, to avoid positioning artifacts described in Text SI and Figure S6. Cropped transects were used in analyses shown in Figure 5 and S7CD. On analyses quantifying differences in intracellular asymmetry, such as in Figure 5, raw measurements were used without the need for length normalization.

Non-normalized transects were also used to obtain the proportion of the mother cell represented by the old area, shown in Figure 6. In this case, each log-transformed transect was fit with a segmented regression using R package “segmented” v.1.4-0 (55). The breakpoint along the cell length marking the transition from the gradient to the fluorescence plateau was used as the old pole length in further analyses.

### Promoter activity quantification

The estimate of RpoS promoter activity was performed as previously described (4, 8). Given the mean fluorescence of a cell (M), its length (L) and a constant (*p* = 0.02), promoter activity (A) is defined as

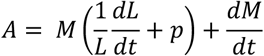

The promoter activity of mother cells over time was performed for all fluorescence measurements throughout the individual’s lifespan. In this case, M was smoothed with a moving window of 5 fluorescence frames, and the resulting A was again averaged over a similar moving window for graphical representation. For the comparison of mother and daughter cells, since the latter was only followed for a generation, measurements (M and L) at the beginning and end of each generation were used to obtain A. No data smoothing was employed in this comparison.

## Data availability

All data and supporting code will be available from a public repository by the time of publication. Strains can be obtained from the Uri Alon library of *E. coli* fluorescent transcriptional reporters (40) and materials will be shared upon request.

## Acknowledgements

We thank the Evolutionary Demography Group members, L. Stäcker and J. Kuom for experimental support, discussions, and feedback. We thank D. Roizman and M. Herzog for kindly providing the constitutive promoter strain. AMP was supported by a Rising Star Postdoctoral Fellowship provided by the FU Berlin and a Humboldt Research Fellowship provided by the Alexander von Humboldt Foundation, Germany. MT was funded by the German Research Foundation (DFG, 430174701) and by the European Union under the Marie SkŁodowska-Curie Actions’ European Postdoctoral Fellowship grant agreement (101069035). US was funded by the German Research Foundation (DFG, 430170797) as a Heisenberg Fellow.

## References

1. A. Urchueguía, et al., Genome-wide gene expression noise in Escherichia coli is condition-dependent and determined by propagation of noise through the regulatory network. PLoS Biol. 19, e3001491 (2021).

2. N. Rosenfeld, J. W. Young, U. Alon, P. S. Swain, M. B. Elowitz, Gene regulation at the singlecell level. Science 307, 1962–5 (2005).

3. C. Wu, et al., Cellular perception of growth rate and the mechanistic origin of bacterial growth law. Proc. Natl. Acad. Sci. U. S. A. 119, 1–9 (2022).

4. O. Patange, et al., Escherichia coli can survive stress by noisy growth modulation. Nat. Commun. 9 (2018).

5. N. Q. Balaban, J. Merrin, R. Chait, L. Kowalik, S. Leibler, Bacterial persistence as a phenotypic switch. Science (80-.). 305, 1622–1625 (2004).

6. O. Gefen, N. Q. Balaban, The importance of being persistent: Heterogeneity of bacterial populations under antibiotic stress. FEMS Microbiol. Rev. 33, 704–717 (2009).

7. P. B. Rainey, et al., The evolutionary emergence of stochastic phenotype switching in bacteria. Microb. Cell Fact. 10, S14 (2011).

8. N. M. V. Sampaio, C. M. Blassick, V. Andreani, J. B. Lugagne, M. J. Dunlop, Dynamic gene expression and growth underlie cell-to-cell heterogeneity in Escherichia coli stress response. Proc. Natl. Acad. Sci. U. S. A. 119, 1–12 (2022).

9. L. Chao, C. U. Rang, A. M. Proenca, J. U. Chao, Asymmetrical Damage Partitioning in Bacteria: A Model for the Evolution of Stochasticity, Determinism, and Genetic Assimilation. PLoS Comput. Biol. 12, e1004700 (2016).

10. A. M. Proenca, C. U. Rang, C. Buetz, C. Shi, L. Chao, Age structure landscapes emerge from the equilibrium between aging and rejuvenation in bacterial populations. Nat. Commun. 9, 3722 (2018).

11. U. K. Steiner, et al., Two stochastic processes shape diverse senescence patterns in a singlecell organism. Evolution (N. Y). 73, 847–857 (2019).

12. S. C. Stearns, The Evolution of Life History Traits: A Critique of the Theory and a Review of the Data. Annu. Rev. Ecol. Syst. 8, 145–171 (1977).

13. M. R. Rose, Evolutionary Biology of Aging (Oxford University Press, 1991).

14. T. B. L. Kirkwood, Evolution of ageing. 270, 301–304 (1977).

15. L. Partridge, N. H. Barton, Optimally, mutation and the evolution of ageing. Nature 362, 305–311 (1993).

16. G. C. Williams, Pleiotropy, Natural Selection and the Evolution of Senescence. Evolution (N. Y). 11, 398–411 (1957).

17. W. D. Hamilton, The moulding of senescence by natural selection. J. Theor. Biol. 12, 12–45 (1966).

18. M. Ackermann, S. C. Stearns, U. Jenal, Senescence in a Bacterium with Asymmetric Division. Science (80-.). 300, 1920 (2003).

19. E. J. Stewart, R. Madden, G. Paul, F. Taddei, Aging and death in an organism that reproduces by morphologically symmetric division. PLoS Biol. 3, e45 (2005).

20. N. Erjavec, M. Cvijovic, E. Klipp, T. Nyström, Selective benefits of damage partitioning in unicellular systems and its effects on aging. Proc. Natl. Acad. Sci. U. S. A. 105, 18764–18769 (2008).

21. S. R. Laney, R. J. Olson, H. M. Sosik, Diatoms favor their younger daughters. Limnol. Oceanogr. 57, 1572–1578 (2012).

22. C. U. Rang, A. Y. Peng, L. Chao, Temporal dynamics of bacterial aging and rejuvenation. Curr. Biol. 21, 1813–6 (2011).

23. L. Jouvet, A. Rodríguez-Rojas, U. K. Steiner, Demographic variability and heterogeneity among individuals within and among clonal bacteria strains. Oikos 127, 0–2 (2018).

24. A. M. Proenca, C. U. Rang, A. Qiu, C. Shi, L. Chao, Cell aging preserves cellular immortality in the presence of lethal levels of damage. PLOS Biol. 17, e3000266 (2019).

25. M. Tuğrul, U. K. Steiner, Demographic Consequences of Damage Dynamics in Single-Cell Ageing. bioRxiv (2023) 10.1101/2023.05.23.538602.

26. S. N. Evans, D. Steinsaltz, Damage segregation at fissioning may increase growth rates: A superprocess model. Theor. Popul. Biol. 71, 473–490 (2007).

27. T. B. L. Kirkwood, Understanding ageing from an evolutionary perspective. J. Intern. Med. 263, 117–127 (2008).

28. L. Chao, A model for damage load and its implications for the evolution of bacterial aging. PLoS Genet. 6, e1001076 (2010).

29. M. Ackermann, L. Chao, C. T. Bergstrom, M. Doebeli, On the evolutionary origin of aging. Aging Cell 6, 235–44 (2007).

30. A. B. Lindner, R. Madden, A. Demarez, E. J. Stewart, F. Taddei, Asymmetric segregation of protein aggregates is associated with cellular aging and rejuvenation. Proc. Natl. Acad. Sci. U. S. A. 105, 3076–81 (2008).

31. A. B. Lindner, A. Demarez, Protein aggregation as a paradigm of aging. Biochim. Biophys. Acta - Gen. Subj. 1790, 980–996 (2009).

32. A. S. Coquel, et al., Localization of Protein Aggregation in Escherichia coli Is Governed by Diffusion and Nucleoid Macromolecular Crowding Effect. PLoS Comput. Biol. 9 (2013).

33. J. Winkler, et al., Quantitative and spatio-temporal features of protein aggregation in Escherichia coli and consequences on protein quality control and cellular ageing. EMBO J. 29, 910–23 (2010).

34. S. K. Govers, J. Mortier, A. Adam, A. Aertsen, Protein aggregates encode epigenetic memory of stressful encounters in individual Escherichia coli cells. PLOS Biol. 16, e2003853 (2018).

35. U. Łapińska, G. Glover, P. Capilla-Lasheras, A. J. Young, S. Pagliara, Bacterial ageing in the absence of external stressors. Philos. Trans. R. Soc. B Biol. Sci. 374 (2019).

36. D. Steinsaltz, M. D. Christodoulou, A. A. Cohen, U. K. Steiner, Chance events in aging (Elsevier Inc., 2019) 10.1016/B978-0-12-801238-3.11394-7.

37. C. Shi, et al., Allocation of gene products to daughter cells is determined by the age of the mother in single Escherichia coli cells. Proc. R. Soc. B 287, 20200569 (2020).

38. O. K. Silander, et al., A genome-wide analysis of promoter-mediated phenotypic noise in Escherichia coli. PLoS Genet. 8 (2012).

39. P. Wang, et al., Robust growth of Escherichia coli. Curr. Biol. 20, 1099–103 (2010).

40. A. Zaslaver, et al., A comprehensive library of fluorescent transcriptional reporters for Escherichia coli. Nat. Methods 3, 623–628 (2006).

41. D.-D. Yang, et al., Fitness and Productivity Increase with Ecotypic Diversity among Escherichia coli Strains That Coevolved in a Simple, Constant Environment. Appl. Environ. Microbiol. 86, e00051–20 (2020).

42. H. Weber, T. Polen, J. Heuveling, V. F. Wendisch, R. Hengge, Genome-wide analysis of the general stress response network in Escherichia coli: sS-dependent genes, promoters, and sigma factor selectivity. J. Bacteriol. 187, 1591–1603 (2005).

43. R. Lange, R. Hengge‐Aronis, Identification of a central regulator of stationary‐phase gene expression in Escherichia coli. Mol. Microbiol. 5, 49–59 (1991).

44. S. Klumpp, Z. Zhang, T. Hwa, Growth Rate-Dependent Global Effects on Gene Expression in Bacteria. Cell 139, 1366–1375 (2009).

45. L. Chao, C. K. Chen, C. Shi, C. U. Rang, Spatial and temporal distribution of ribosomes in single cells reveals aging differences between old and new daughters of Escherichia coli. Elife 12, RP89543 (2023).

46. S. Taheri-Araghi, et al., Cell-Size Control and Homeostasis in Bacteria. Curr. Biol. 25, 385–391 (2015).

47. Y. Yang, et al., Temporal scaling of ageing as an adaptive strategy of Escherichia coli. Sci. Adv. 5, eaaw2069 (2019).

48. L. K. Poulsen, T. R. Licht, C. Rang, K. A. Krogfelt, S. Molin, Physiological state of Escherichia coli BJ4 growing in the large intestines of streptomycin-treated mice. J. Bacteriol. 177, 5840–5845 (1995).

49. J. Mortier, et al., Protein aggregates act as a deterministic disruptor during bacterial cell size homeostasis. Cell. Mol. Life Sci. 80, 1–13 (2023).

50. L. Wolf, O. K. Silander, E. van Nimwegen, Expression noise facilitates the evolution of gene regulation. Elife 4, e05856 (2015).

51. J. B. Lugagne, H. Lin, M. J. Dunlop, DeLTA: Automated cell segmentation, tracking, and lineage reconstruction using deep learning. PLoS Comput. Biol. 16, 1–18 (2020).

52. R Core Team, R: A language and environment for statistical computing (2017).

53. S. N. Wood, Fast stable restricted maximum likelihood and marginal likelihood estimation of semiparametric generalized linear models. J. R. Stat. Soc. Ser. B Stat. Methodol. 73, 3–36 (2011).

54. K. P. Burnham, D. R. Anderson, Multimodel inference: Understanding AIC and BIC in model selection. Sociol. Methods Res. 33, 261–304 (2004).

55. V. M. R. Muggeo, Estimating regression models with unknown break-points. Stat. Med. 22, 3055–3071 (2003).

