## Supplementary Materials for "Declining old pole physiology gradually enhances gene expression asymmetry in bacteria"

**This PDF file includes:**

SI Text
Figures S1 to S6
Tables S1 to S3

**Supporting Information Text**

**Controlling for positioning artifacts.** The mother machine was designed to retain the mother cell at the closed end of growth wells (Figure 1A). Consequently, its old pole has no neighboring cells, while its new pole has one neighbor. The daughter cell, as shown in the time-lapse images in Figure 1B and 1F, usually has neighbors on both ends. Because some of the fluorescence expressed by each individual might scatter over its neighbors, it is possible that the daughter cell appears brighter than the mother due to this neighboring effect. Therefore, to confirm that the mother-daughter asymmetry reported in Figure 2 has physiological origins, we performed additional control analyses.

As a control for differences in mean fluorescence, we looked at the first cell division observed after cells were loaded into the device (Figure S3). The polarity of the founder cell is unknown. We can assume either pole could be at the closed end early on, with equal probability. Thus, upon the first division, the "2nd cell" (as labeled in Figure S3A) might inherit either a new or an old pole from the "1st cell". We hypothesize that the fluorescence asymmetry reported in Figure 2 is correlated with pole inheritance, which means that we expect to observe no clear asymmetry between 1st and 2nd cells due to their mixed polarity. If we were to find asymmetry at this early division, it would mean that the patterns here reported are merely an artifact of a cell's positioning within the mother machine. By performing these measurements (Figure S3B-D), we observed no difference in mean fluorescence between 1st and 2nd cells. After the following division, we know that the cell by the closed end has retained an old pole, and that the other cell received a new pole (Figure S3A), a pattern that will remain unchanged for the rest of the experiment. From this point onwards, we observed a growing asymmetry between mother and daughter cells (Figure S3E-F). We also verified that this asymmetry was not a function of the physical proximity between mothers and daughters (Figure S4), with maternal intracellular fluorescence ratios < 1 even when the daughter was positioned further away in the growth well. These results indicate that differences in mean fluorescence are not a positioning artifact, but a pattern that appears once subpopulations of mothers and daughters are defined by cell pole inheritance.

Even after verifying that asymmetry has a physiological origin, however, we must consider that positioning artifacts might still impact our measurements. This becomes more relevant as we move from average measurements to the quantification of intracellular gradients, such as shown in Figure 4. The gradients exhibited by mother cells could be due to the absence of neighbors on the old pole side ("outer pole" in Figure S5A), while its new pole receives scattered fluorescence from the daughter cell. To determine whether this was the case, we estimated the intracellular gradients of cells by the open end of the growth well. These cells also have neighbors on only one side, and their polarity can be determined according to the number of cells in the well at a given moment (Figure S5A). When there are either 3, 7, or 11 cells in a well, the outermost cell is likely a new-pole daughter, with its new pole facing outwards ("outer pole"). By measuring the intracellular fluorescence of the outer cells (Figure S5B-C) we observed that, although outer poles are generally darker than inner poles, the maternal outer pole was darker and its gradient more pronounced. While the intracellular asymmetry of mother cells increased with consecutive old pole inheritances (Figure S5C), the outer cell became more symmetric over time. Our expectation that this outer cell would exhibit intracellular symmetry, however, was not met (Figure S5D), indicating that the fluorescence scattered by neighboring cells influences gene expression gradients.

To address this issue, we measured the fluorescence scattered by new poles (Figure S5A). Following the approach described above to locate daughter cells with new poles facing outwards, we quantified the decline in background fluorescence as a function of distance from the outer cell (Figure S5E). Since this outer pole is equivalent to the new pole neighboring the mother cell, this provides a good estimate of the light scattered over the mother. We found a significant “scatter zone” stretching over the first 10 pixels, after which fluorescence signals were indistinguishable from true background noise (Figure S5F). To avoid biases produced by this scatter, we measured intracellular gradients after cropping 10 px on each side of all cells (Figure 4). This was a conservative choice, without considering the gaps between cells (shown in Figure S4). As the resulting transects in Figure 4 show, the mother cell retained its intracellular asymmetry, and the daughter its symmetry.

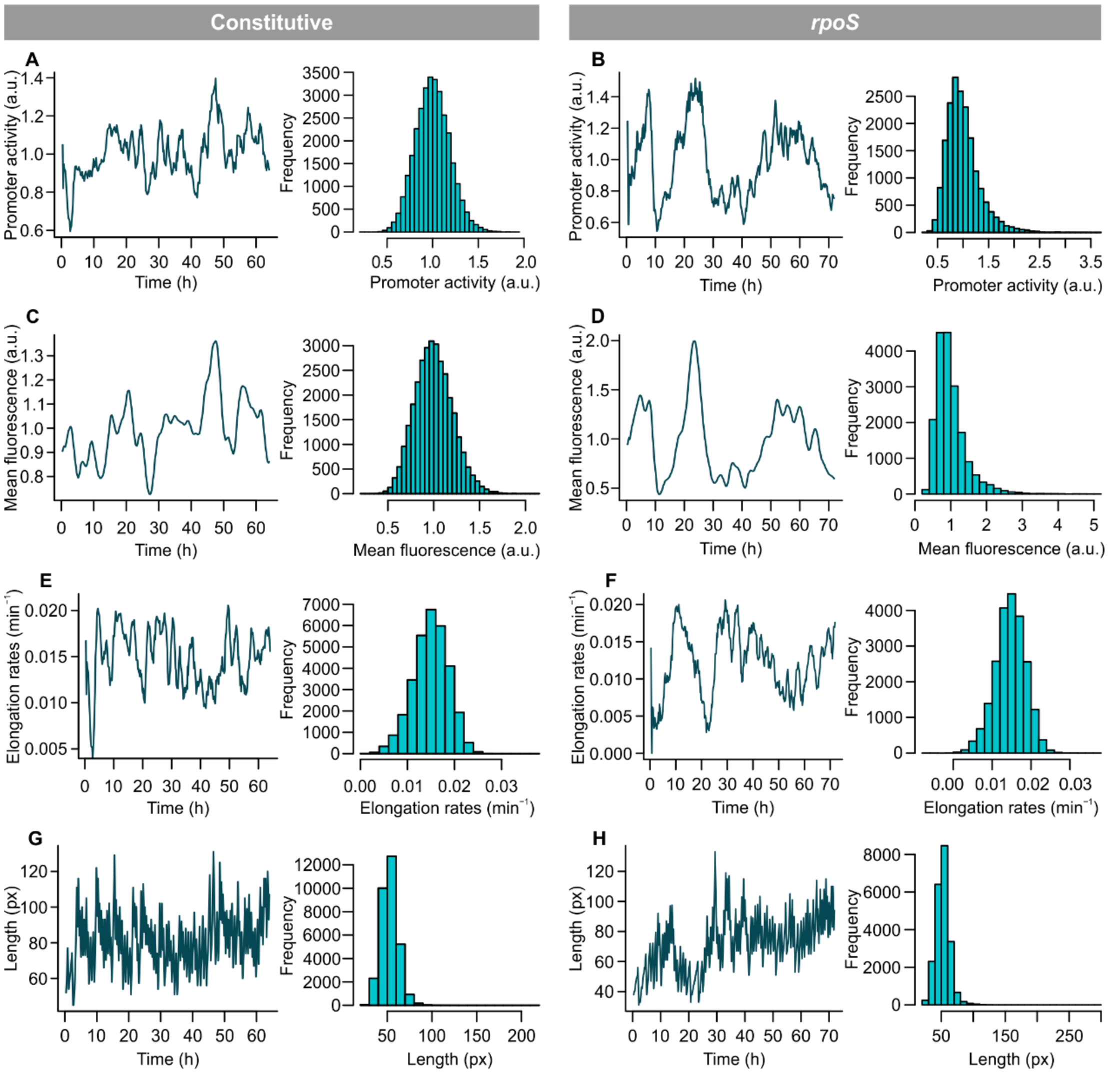


**Fig. S1. GFP expression and growth physiology.** Panels on the left show a representative mother cell lineage over up to 100 generations. Panels on the right show histograms for all mother cell lineages, considering one measurement per generation. ($n_{Const}$ = 31,472, $n_{rpoS}$ = 18,425). (A and B) Promoter activity calculated from cell lengths, elongation rates, and mean fluorescence. While values on the left were calculated at each time point (10 min intervals), values in histograms were simplified to consider one estimate per generation. This estimate is also used in Figures 1 and 3 (see Methods for details). (C and D) Mean fluorescence levels, showing fluctuations over time. While *rpoS* shows higher fluctuations and a long-tailed distribution, constitutive GFP expression has lower noise levels. (E and F) While no correlation is observed between constitutive GFP levels and fluctuations in elongation rates, *rpoS* fluorescence showed a negative correlation with growth. (G and H) Cell length over time. Peaks of RpoS fluorescence also corresponded to smaller cell lengths, because of growth inhibition.

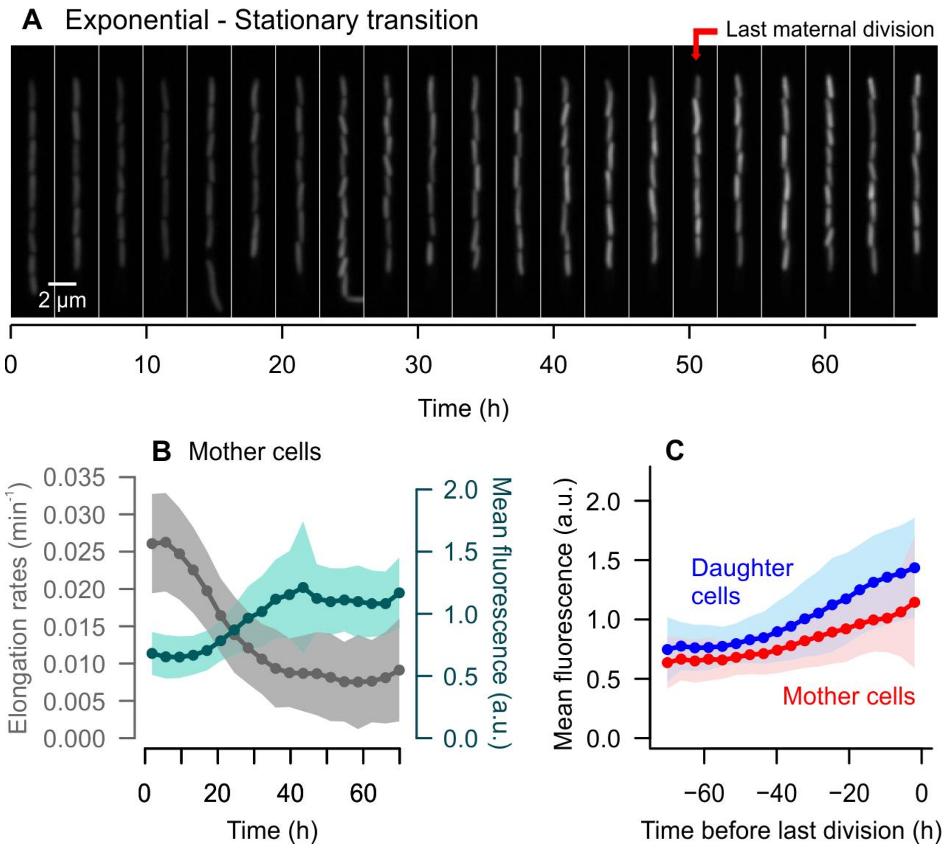


**Fig. S2. RpoS activity increases as cells enter stationary phase.** (A) Time-lapse of GFP expression, showing a mother cell (on top) and its daughters over time. After ~50 h, the mother cell switches from an active to a stationary growth state, while its daughters remain active. The image shows a gradual increase in fluorescence, which is summarized in (B). The average elongation rates of the population comprised by mother cells declined over time, as the growth medium was replaced with spent medium. RpoS fluorescence increased, in agreement with the growth-inhibitory activity of this transcription factor in the transition from exponential to stationary phase. (C) If we align the timeline of all lineages by the last maternal division (as shown by the arrow in A), we observe an increase in RpoS fluorescence as each lineage reaches stationary phase. A generalized additive model on log-transformed fluorescence measurements ($n = 64{,}947$) indicates that fluorescence varied smoothly over time, with different curves representing mother ($F = 3{,}758$, $p < 0.001$) and daughter cells ($F = 6{,}562$, $p < 0.001$). A cell's pole age (new or old pole inheritance) also impacted mean fluorescence levels as a linear predictor ($t = 123.6$, $p < 0.001$). Together, the model given by ln(GFP) ~ s(time, by = Age) + Age captured 61.9% of the variance observed in RpoS fluorescence during the transition to stationary phase. (B and C) Bins = means ± SD.

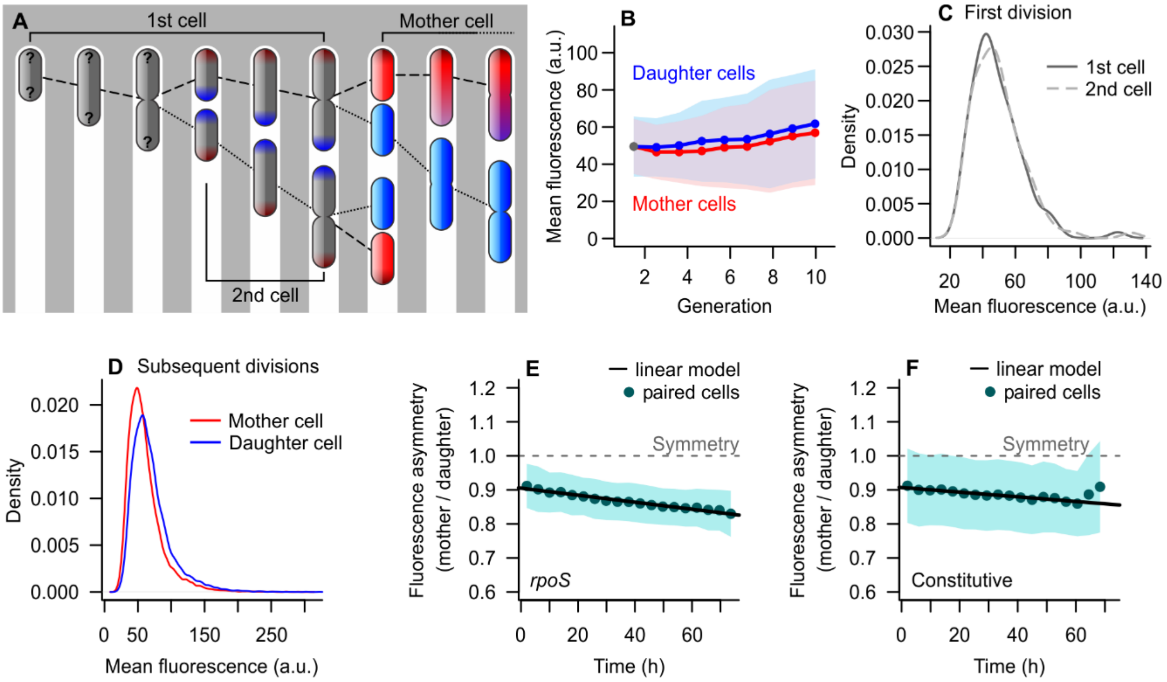


**Fig. S3. Mean fluorescence asymmetry is not an artifact of mother machine positioning biases.** Since mother cells are always positioned at the closed edge of the growth wells and daughter cells occupy the middle of such traps, we performed control analyses to ensure that the asymmetry observed in Figure 2C to 2F does not arise from an imaging artifact. (A) We looked at the early divisions, shortly after the cells were loaded into the device. Since the polarity of the first cell is unknown, it is not possible to determine which cell inherits old or new poles upon the first division. As such, we expect no clear asymmetry between the 1st and 2nd cells. (B) By looking at the mean fluorescence after each division, we observe that the asymmetry only begins to appear after some divisions and once the distinction between mother and daughter cells is determined. The first division is represented by gray dots. The distributions of mean fluorescence measurements for these points are shown in (C), indicating no asymmetry between 1st and 2nd cells. By comparing these distributions to those of the entire population across time (D), we can conclude that the asymmetry observed in Figure 2 has physiological origins. (E and F) This is further supported by the fact that mother-daughter asymmetry increases as mother cells age, which was verified for both promoters (*rpoS*: $t = -48.58$, $p < 0.001$; Constitutive: $t = -20.97$, $p < 0.001$).

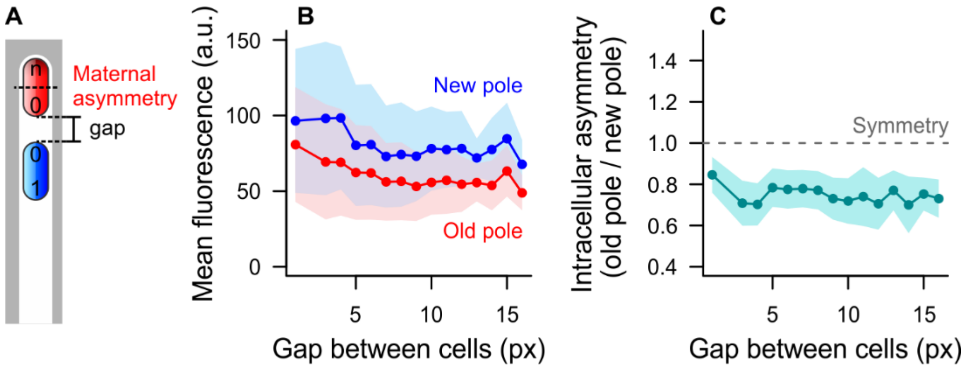


**Fig S4. Controls for neighbor artifacts on mean fluorescence measurements.** As explained in SI Text, mother cells receive some light scattered from the new pole of their daughters, which can create confounding results. (A) To determine whether the intracellular asymmetry of mother cells was produced by the proximity to its daughter, we measured the spatial gap between mothers and daughters. The first 5 h of growth were not considered, as the effects of asymmetry take a few generations to stabilize. (B) Maternal intracellular asymmetry was present independently of the gap (Wilcoxon signed rank test, $V = 2108840$, $p < 0.001$). Fluorescence measurements were higher when the daughter was closer, stabilizing for gaps > 10 pixels. (C) The ratios between maternal old and new poles were not largely affected. This indicates that asymmetry was present despite any positioning artifacts and neighbor light scattering. Bins = mean ± SD.

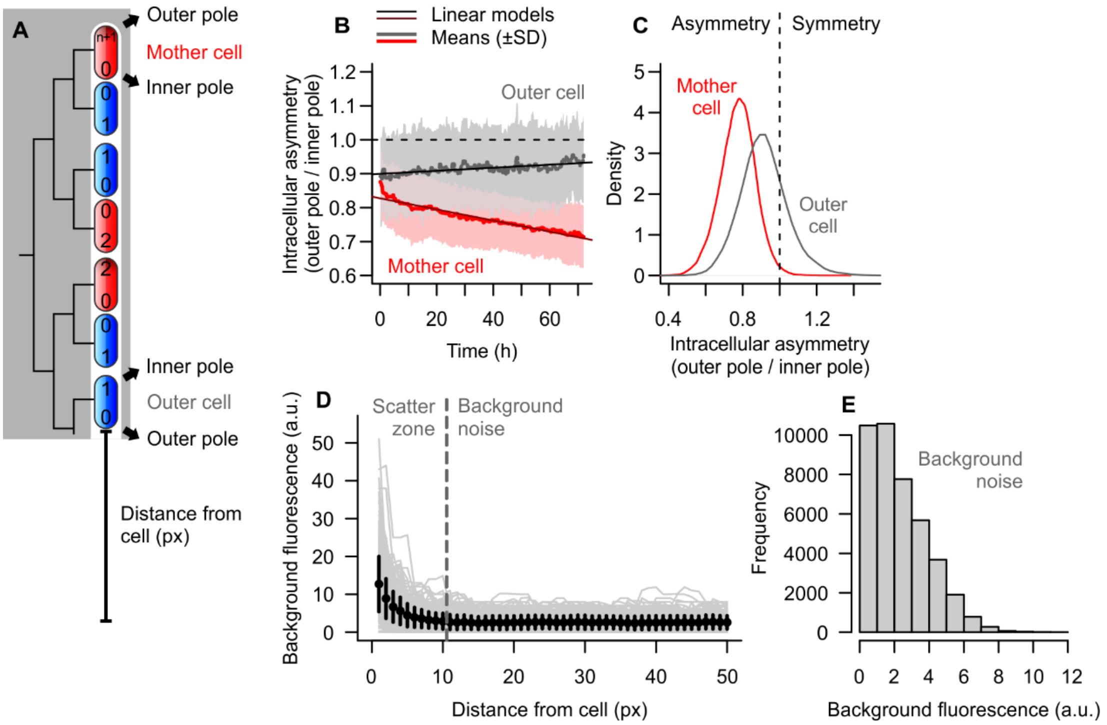


**Fig. S5. Positioning artifacts on fluorescence measurements.** (A) The mother machine was designed to retain the mother cell at the closed end of growth wells. Consequently, its old pole (age = n+1) has no neighboring cells, while its new pole (age = 0) has one neighbor. This new pole might receive fluorescence scattered by the neighboring cell, introducing measurement biases (see SI Text). (B and C) Dashed line indicates gene expression symmetry. (B) By comparing outer and inner poles, we observed that the maternal intracellular asymmetry increased over time ($\beta = -0.011$, $t = -110.2$, $p < 0.001$; $n = 90{,}277$), while outer cells had a slight increase in symmetry ($\beta = 0.0005$, $t = 8.312$, $p < 0.001$; $n = 26{,}356$). (C) Overall, both mothers (ratio = 0.773) and outer cells (ratio = 0.915) were significantly asymmetric ($t_{mother} = -709.6$, $t_{outer} = -77.5$, $p < 0.001$). Since outer cells could be expected to exhibit symmetry comparable to the daughters, this would mean that neighboring effects produce an artificial ~8.5% asymmetry. Thus, we corrected the fluorescence transects in Figure 4 as follows: (D) To quantify the fluorescence scattered by the daughter's new pole over mother cells, we looked for poles of equivalent age located at the open edge of growth wells. We measured the decay in fluorescence signals as the distance from the cell increased, finding that there is a significant scatter over the first 10 pixels (Wilcoxon rank sum test with Bonferroni correction, $p < 0.001$ for px = 1 to 10). Bins = mean ± SD. (E) Fluorescence measurements obtained >50 pixels away from any cells show the distribution of true background values, used as a null distribution for tests in (D).

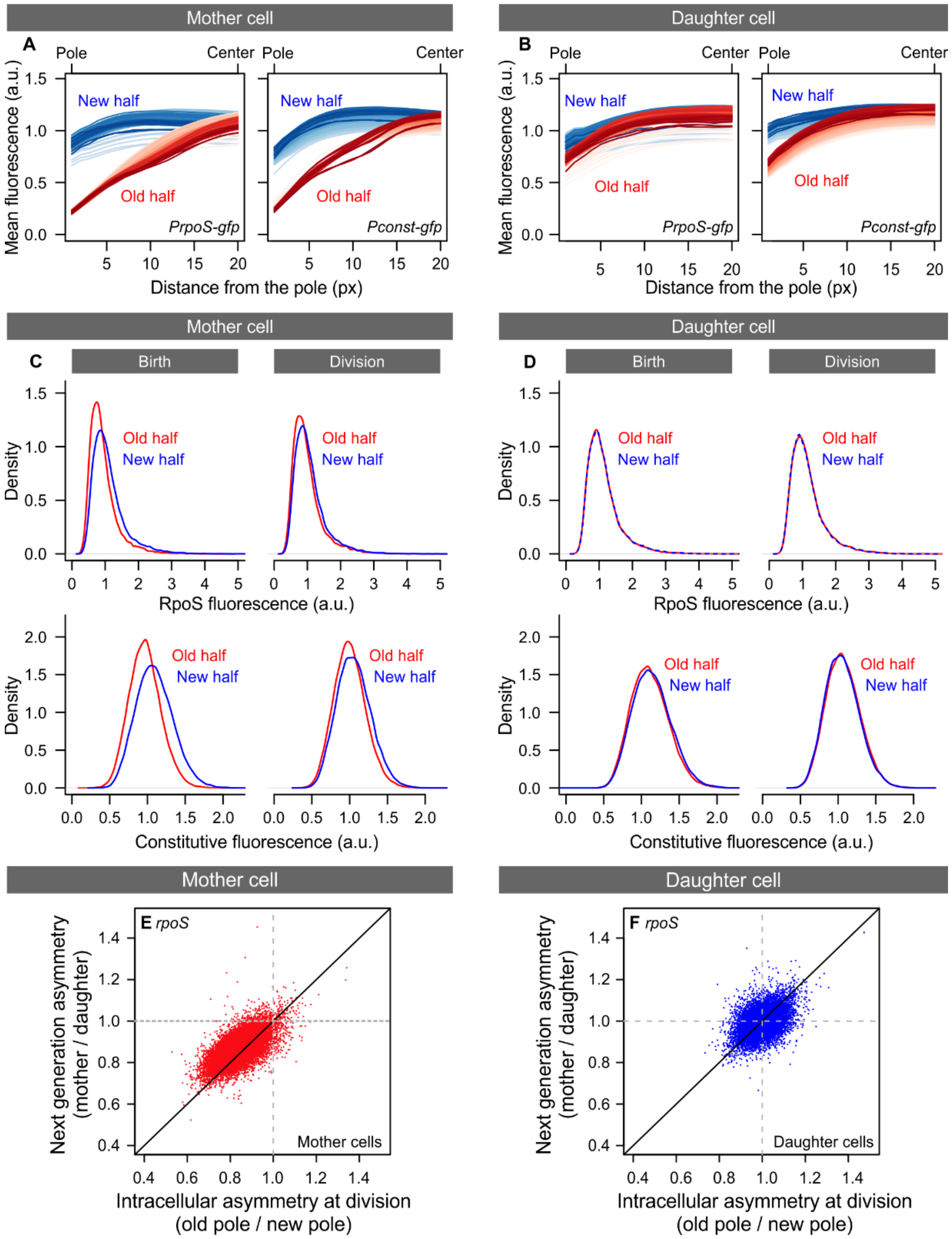


**Fig. S6. Intracellular asymmetry in gene expression.** (A and B) Folded representation of transects from Figure 4, as used for fitting generalized linear models. Rather than considering the fluorescence across the entire length of the cell, the distance from the pole towards the center was used as a smoothing term (see also Table S2). Each transect represents an average of all cells in a given generation, with darker line colors representing later generations. (C) Normalized fluorescence distributions of old and new cell halves, estimated from cropped transects in Figure 3, show that mother cells exhibit intracellular asymmetry after correcting for positioning artifacts. At birth, the maternal old halves showed higher *rpoS* fluorescence than new halves (paired t test, $t = 222.63$, $p < 0.001$; $n = 18{,}979$ cells), which was also the case for constitutive expression ($t = 197.75$, $p < 0.001$; $n = 32{,}399$ cells). This asymmetry decreased throughout the cell cycle, but remained significant by the time of division (*rpoS*: $t = 149.98$, $p < 0.001$; constitutive*:* $t = 93.21$, $p < 0.001$). (D) Daughter cells, on the other hand, were nearly symmetric, although statistical analysis still captured a significant difference between cell halves for both *rpoS* ($t = 19.77$, $p < 0.001$; $n = 18{,}444$) and the constitutive promoter ($t = 61.85$, p <

0.001; $n = 31{,}258$). (E and F) The intracellular asymmetry that a cell displayed before division was positively correlated with the asymmetry among its offspring after division. There was a significant correlation for both mother cells (Pearson's $r = 0.658$; $t = 118.740$, $p < 0.001$) and daughter cells (Pearson's $r = 0.454$; $t = 64.599$, $p < 0.001$).

**Table S1.** Variance partitioning of fluorescence measurements presented in Figure 2, performed on log-transformed data. An interaction term between elongation rates and age was not significant for the constitutive promoter.

***rpoS***

| | Sum Sq | Var explained [95% CI] | | F | p |
|---|---|---|---|---|---|
| Elongation rates | 77.653 | 23.496 % | [22.639 - 24.323 %] | 7971.638 | < 0.001 |
| Age (mother or daughter) | 32.606 | 9.866 % | [9.378 - 10.369 %] | 6694.534 | < 0.001 |
| Maternal lineage | 38.055 | 11.515 % | [11.358 - 12.413 %] | 32.154 | < 0.001 |
| Time | 2.769 | 0.838 % | [0.704 - 0.973 %] | 568.575 | < 0.001 |
| Elongation rates x Age | 0.787 | 0.238 % | [0.169 - 0.312 %] | 80.779 | < 0.001 |
| Residuals | 178.628 | 54.048 % | [52.849 - 54.565 %] | - | - |

Model: ln(Mean fluorescence) ~ poly(Elongation rates,2) * Age + Lineage + Time

**Constitutive**

| | Sum Sq | Var explained [95% CI] | | F | p |
|---|---|---|---|---|---|
| Elongation rates | 0.769 | 1.556% | [1.371 - 1.749 %] | 892.630 | < 0.001 |
| Age (mother or daughter) | 5.848 | 11.829% | [11.346 - 12.322 %] | 13572.220 | < 0.001 |
| Maternal lineage | 12.393 | 25.067% | [24.942 - 26.017 %] | 65.666 | < 0.001 |
| Time | 3.675 | 7.434% | [7.075 - 7.696 %] | 8529.339 | < 0.001 |
| Residuals | 26.753 | 54.114% | [53.233 - 54.300 %] | - | - |

Model: ln(Mean fluorescence) ~ poly(Elongation rates,2) + Age + Lineage + Time

**Table S2.** Generalized additive model selection for mother cells shown in Figure S7A and S7B. The models consider the variation in gene expression across the cell length, from each pole towards the center (*dist*), its changes over *time* and differences among new and old halves of the cell (*pole*). The highlighted model exhibited the lowest AIC and explained the largest fraction of the deviance in our data. Fluorescence measurements were log-transformed prior to model fitting, without cropping. See Table S3 for statistical output.

***rpoS***

| Model | df | AIC | AICΔ | Dev. Exp. |
|---|---|---|---|---|
| **ti(dist,time) + s(dist,by=pole) + s(time,by=pole) + pole** | **50.610** | **760388** | **0** | **51.30%** |
| ti(dist,time) + s(dist,by=pole) + s(time) + pole | 41.511 | 763328 | 2941 | 51.10% |
| ti(dist,time) + s(dist) + s(time,by=pole) + pole | 42.725 | 934978 | 174590 | 38.50% |
| ti(dist,time) + s(dist) + s(time) + pole | 33.815 | 937305 | 176917 | 38.30% |
| ti(dist,time) + s(dist,by=pole) + s(time,by=pole) | 48.457 | 960458 | 200070 | 36.40% |
| ti(dist,time) + s(dist) + s(time) | 32.372 | 1099505 | 339117 | 23.40% |

**Constitutive**

| Model | df | AIC | AICΔ | Dev.Exp. |
|---|---|---|---|---|
| **ti(dist,time) + s(dist,by=pole) + s(time,by=pole) + pole** | 54.514 | 124904 | 0 | **67.10%** |
| ti(dist,time) + s(dist,by=pole) + s(time) + pole | 45.422 | 138847 | 13943 | 66.80% |
| ti(dist,time) + s(dist) + s(time,by=pole) + pole | 45.329 | 572958 | 448054 | 53.60% |
| ti(dist,time) + s(dist) + s(time) + pole | 36.505 | 582836 | 457932 | 53.20% |
| ti(dist,time) + s(dist,by=pole) + s(time,by=pole) | 53.074 | 591878 | 466974 | 52.30% |
| ti(dist,time) + s(dist) + s(time) | 35.553 | 927491 | 802587 | 39.00% |

**Table S3.** Statistically significant predictors of the model highlighted in Table S2, applied to mother and daughter cell fluorescence transects. The final model had a significant tensor product interaction between distance and time, and a significant difference between new and old cell halves. The individual smoothing terms varied according to cell half, with both distance and time exhibiting a significant effect on gene expression. Models that did not distinguish new and old cell halves had worse fit to the data (Table S2).

***rpoS***

| | Mother cell | | | Daughter cell | | |
|---|---|---|---|---|---|---|
| **Linear predictor** | **Estimate** | **t** | **p-value** | **Estimate** | **t** | **p-value** |
| Pole | -0.445 | -478.9 | <0.001 | -0.049 | -54.57 | <0.001 |
| **Non-linear predictor** | **Edf** | **F** | **p-value** | **edf** | **F** | **p-value** |
| ti(dist,time) | 13.576 | 70.08 | <0.001 | 7.843 | 12.53 | <0.001 |
| s(dist):new half | 7.138 | 1,517.6 | <0.001 | 8.08 | 2,123.25 | <0.001 |
| s(dist):old half | 8.975 | 59,429.02 | <0.001 | 7.979 | 6,390.73 | <0.001 |
| s(time):new half | 8.981 | 952.79 | <0.001 | 8.942 | 999.38 | <0.001 |
| s(time):old half | 8.941 | 356.88 | <0.001 | 8.861 | 1,375.63 | <0.001 |
| **Deviance explained** | | **51.3%** | | | **11.7%** | |

**Constitutive**

| | Mother cell | | | Daughter cell | | |
|---|---|---|---|---|---|---|
| **Linear predictor** | **Estimate** | **t** | **p-value** | **Estimate** | **t** | **p-value** |
| Pole | -0.335 | -749.8 | <0.001 | -0.114 | -291.0 | <0.001 |
| **Non-linear predictor** | **Edf** | **F** | **p-value** | **edf** | **F** | **p-value** |
| ti(dist,time) | 15.788 | 362.6 | <0.001 | 8.842 | 26.98 | <0.001 |
| s(dist):new half | 8.917 | 14,390.1 | <0.001 | 8.882 | 3,161.11 | <0.001 |
| s(dist):old half | 8.982 | 211,926.0 | <0.001 | 8.957 | 45,804.8 | <0.001 |
| s(time):new half | 8.834 | 4,563.0 | <0.001 | 8.903 | 44,61.04 | <0.001 |
| s(time):old half | 8.992 | 291.9 | <0.001 | 8.973 | 6,434.16 | <0.001 |
| **Deviance explained** | | **67.1%** | | | **33.3%** | |
